# High concentrations of boric acid induce autophagy in cancer cell lines

**DOI:** 10.1101/193441

**Authors:** Ruslan Al-Ali, Rogelio Gonzalez-Sarmiento

**Affiliations:** Molecular Medicine Unit -IBSAL, Departament of Medicine, University of Salamanca-University hospital of Salamanca-CSIC, Salamanca, Spain; Institute of Molecular and Celular Biology of Cancer (IBMCC), University of Salamanca-CSIC, Salamanca, Spain

**Keywords:** Boric acid, autophagy, cancer, Boron, chloroquine

## Abstract

**Background/Aim:** Boric acid (BA) is thought to have anticancer effects, but only a handful of studies tackled this subject. Though a very common compound, little is known about its therapeutic value, mechanisms and effective doses. This study investigates into its therapeutic value and autophagy as a possible mechanism.

**Materials and Methods:** We evaluated the potency of BA treatment in seven different cell lines. We hypothesized that autophagy is involved in the mechanism of BA toxicity in tumor cells based on observations in plants, insects and cancer cell lines. Changes in autophagy-related proteins were measured after BA treatment. Finally, we suspected that blockage of autophagy would increase the effectiveness of BA treatment and enable the use of smaller doses.

**Results:** Our results demonstrate that all studied cell lines did not suffer mortality in low to medium doses of BA (up to 5mM). However, a high dose (over 25mM) could inflict significant death in all cell lines. Those high doses caused P62/SQSTM1 consumption and LC3II-B accumulation after 3 days of treatment. Using small doses of BA in combination with autophagy blockage did not improve cytotoxicity in lung cancer cell lines.

**Conclusion:** We conclude that high concentrations of BA affect autophagy in short-term treatments. Not enough data is available about BA toxicity, so BA use as cancer treatment can be possible if new toxicity studies are performed.

## 1. Introduction

Boron is an essential trace element for human health. Boron seems to affect the way the body handles other minerals such as magnesium and calcium [1]. It increases estrogen and β-estradiol levels in post-menopausal women so it might act as hormone replacement therapy [2].

The most common inorganic form of Boron is Borax or Boric Acid (BA). Several studies suggested that boric acid had anti-tumoral properties [3, 4, 5, 6]. The mechanism of its lethal effects did not involve apoptosis, but it is suspected to be through histone deacetylase inhibition [7, 8]. Furthermore, boric acid toxicity was reported to be associated with autophagy in plants and insects [9, 10]. While on the other hand, several studies linked histone deacetylase inhibitors (HDACi) with an increased autophagy [11]. Those two facts raised the question of potential role of autophagy in BA mechanism of cytotoxicity, leading to this article.

Toxicity with boric acid in animals causes infertility, neural symptoms and skin irritation. It was reported that children and infants are particularly susceptible to Boron (boric acid) toxicity. However, no sufficient studies *in vivo* are available that could confirm boron/boric acid toxicity or confirm its usefulness as anti-cancer agent. [12]

In this article, we aim to investigate the potency of boric acid as anti-tumoral agent and its effects on autophagy pathway. We use several concentrations of boric acid with different cell lines, and analyse some proteins that are related to autophagy.

## 2. Material and methods

### 2.1. Cell lines

We used seven different cell lines H1299 and COR-L23 (non-small cell lung cancer cell lines), HT-29 and HCT116 (colon), MFC7 and HCC1937 (breast cancer) and CAL-33 (tongue). Cells were cultured in the medium recommended by manufacturer (Sigma-Aldrich), 10% fetal bovine serum (Gibco) and 1% antibiotic (Gibco).

### 2.2. Cell growth

Cell growth was measured using MTT protocol after one week of cell culture (CO2 5% at 37°C). Samples were used in triplets, incubated with MTT (Sigma) for an hour and the dye dissolved in DMSO (Merk Millipore). The measurements were performed via TECAN Infinite^®^ F500. The results were normalized using untreated control.

Cells were treated with BA and/or chloroquine when reached 70-90% of confluency.

### 2.3. Western-blot

Primary anti-rabbit antibodies for P62/SQSTM1 (Abcam: ab109012) and LC3B (NOVUS: NB600-1384) were used, a primary anti-mouse beta-actin antibody (Sigma: A5441), a secondary HRP anti-anti-rabbit (Merck: AP307P) and a secondary HRP anti-anti-mouse (GE: NXA931).

SDS-PAGE gel 8% and 12% and transferred to PVDF membranes (GE) using BioRad electrophoresis and transfer instruments.

## 3. Results

We investigated the use of BA in seven cell lines with different origins. We found that response to BA was dose related, and the LD50 after one-week treatment with BA ranged from 10 to 50mM. We found that 5mM (about 700 folds the normal serum value [13]) affects the growth of the cell lines less than 25%, while only very high concentrations (≥25 mM) cause a significant decrease in cell numbers after a week. (Figure 1)

**Figure 1:**
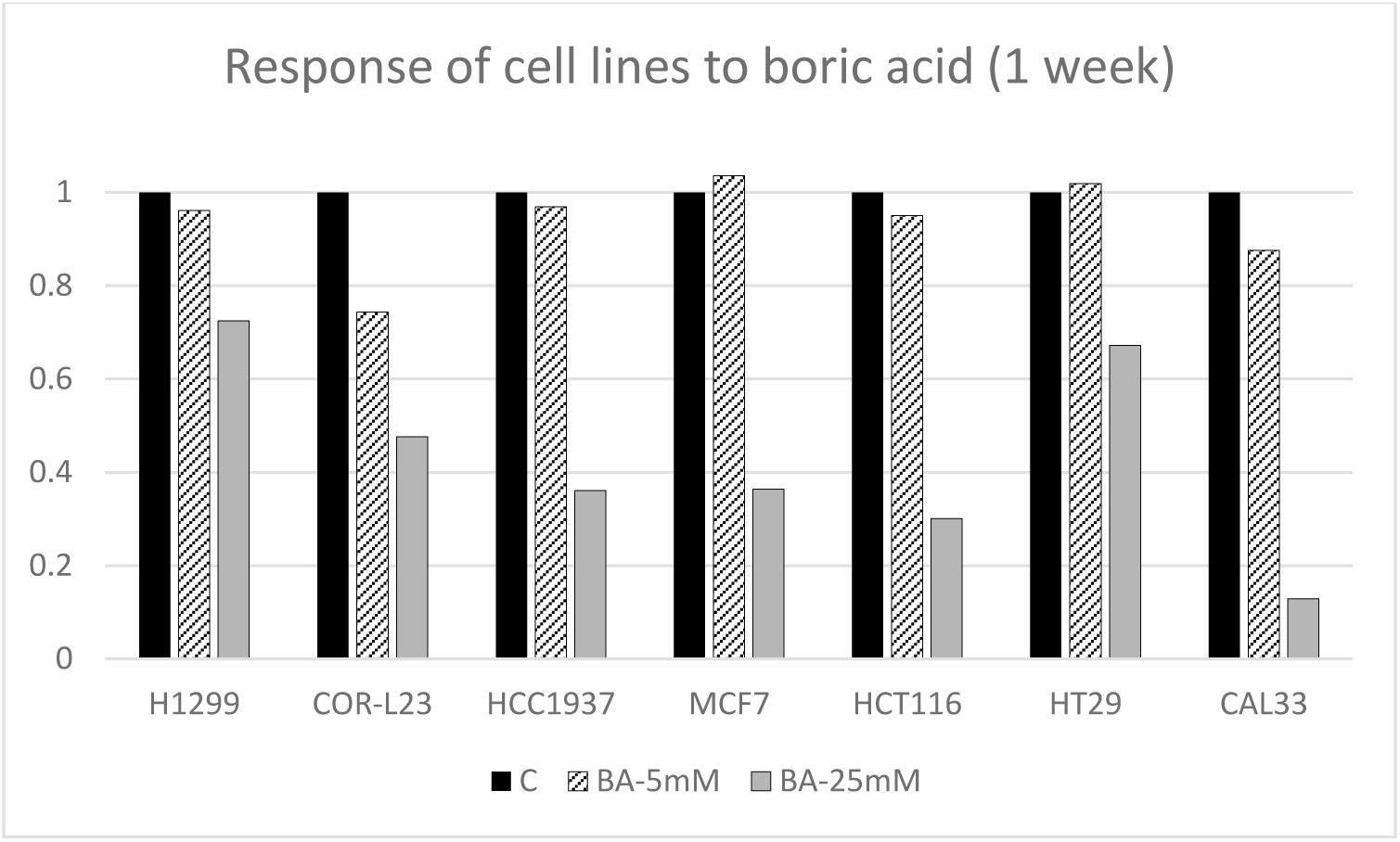
MTT results (normalized) after a week of boric acid treatment for several cell lines with two concentrations: 5mM and 25mM. Cells show limited decrease in growth with 5mM compared to control, but the decrease was stronger with 25mM in all cell lines.

**Figure 2:**
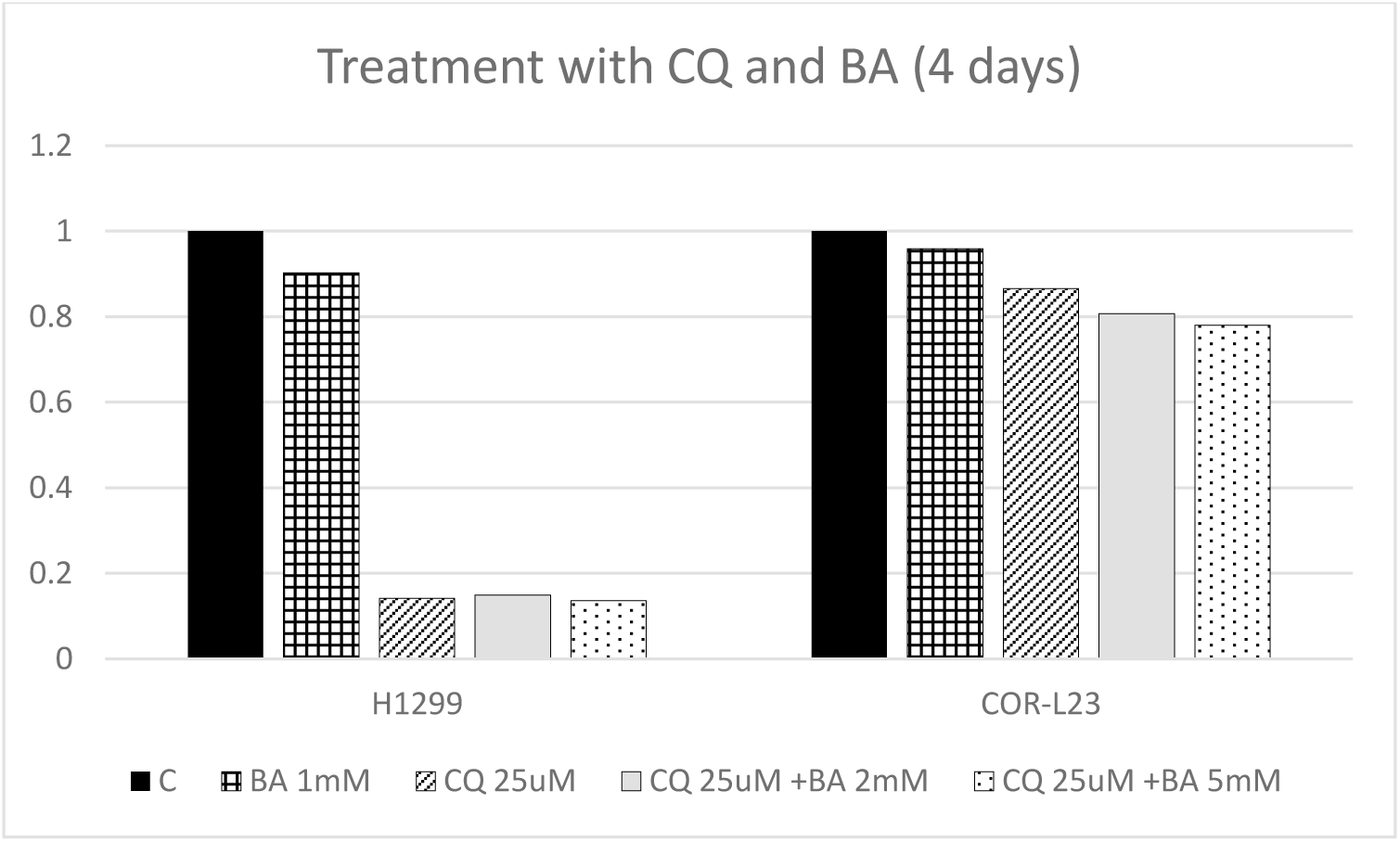
MTT results (normalized) after 4 days with different concentration chloroquine and BA. No significant change between using chloroquine alone and combining it with BA.

To evaluate the relationship between BA toxicity and autophagy, we used H1299 and COR-L23p, two non-small cell lung cancer cell lines (NSCLC), to measure P62 and LC3B after treatment with 5mM and 25mM of BA for 1 week. At 5mM no changes were detected in Western-blot (Not shown). However, with 25Mm of BA, levels of P62/SQSTM1 increased in the first two days then decreased steadily, while a decrease was noticed in LC3B in the first 24 hours then an increase of LC3B-II was detected. (Figure 3 and 4)

**Figure 3:**
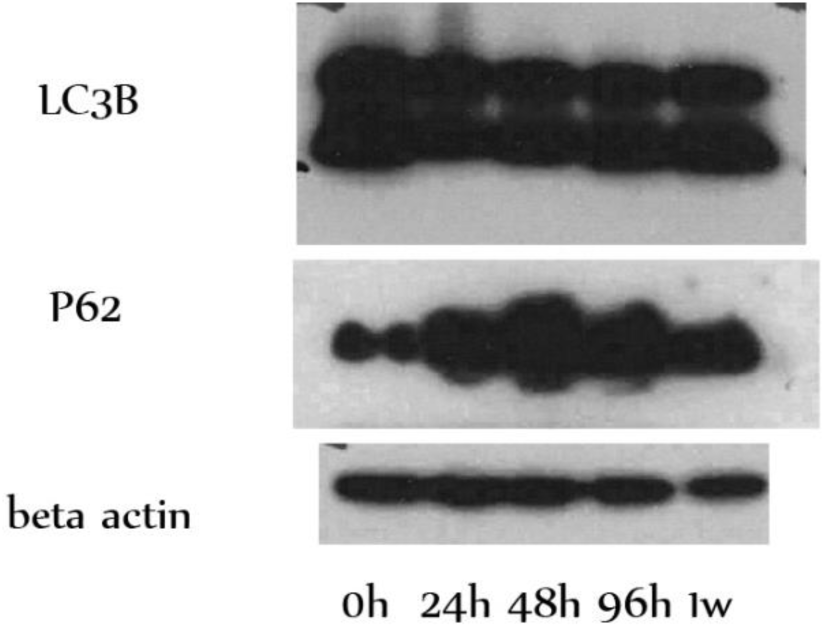
H1299 cell line treated with 25mM of BA. P62 increases for 48h then decreases gradually. LC3B decreases at 24h then LC3B-II increases gradually.

**Figure 4:**
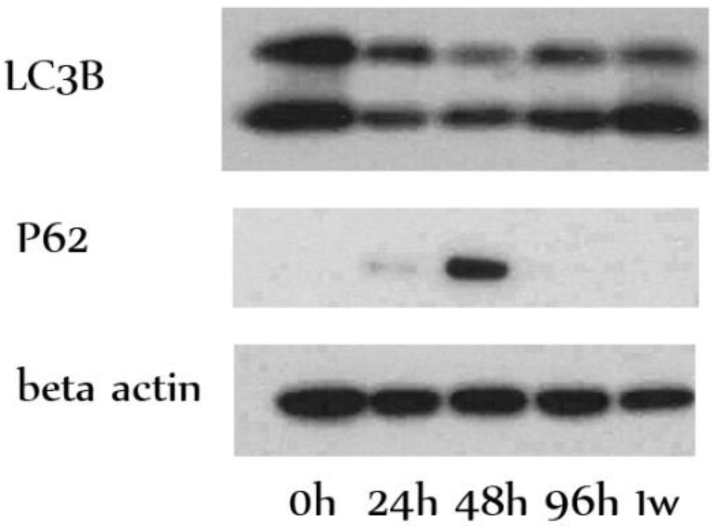
COR-L23p cell line treated with 25mM of BA. P62 increases for 48h then decreases gradually. LC3B decreases at 24h then LC3B-II increases gradually.

The effects on autophagy could be a key for the differences in BA lethality among cell lines, so we studied the effects of BA treatment when the autophagy pathway is blocked using 25µM of chloroquine. In the same NSCLC cell lines, we found that the differences between chloroquine treatment alone for four days and combination with BA are resulting in small changes with 2mM of BA (+5% to −7%), and small decreases with 5mM of BA (−2.5 to −10%). None were statistically significant (p>0.05). (Figure 2)

## 4. Discussion and Conclusions

High nutritional boron intake was suggested to be a protector against lung cancer [14]. Boron is supplied in the diet especially vegetables and fruits, and elevated levels of boron/boric acid exist in several natural spring’s waters that might explain their “miracles properties” [12, 15]. The tolerable upper intake level (UL) for adults over 19 years is 20 mg/day of boron [16]. It was estimated that supplying Boron as boric acid might be toxic with minimal lethal dose for adults is 10-20g [12]. However, cases of boric acid poisoning (10-88 g) reported no fatalities and 88% of them were asymptomatic [12]. The uncertainty about the toxicity led us to explore BA treatment at a very high concentration.

Several studies reported a response in one cell line of prostate cancer (DU-145) to one week of boric acid therapy at 1mM [3, 4, 17]. This cell line is also known as an example of an autophagy-defective cell that lacks *ATG5* [18, 19], while other cells-lines with normal autophagy function do not have a similar response to the same treatment. Boric acid, while affects growth, did not induce apoptosis in breast cancer cells [7]. Studies in flora and insects already pointed to autophagy elevation in boric acid toxicity [9, 10], which was behind the reasoning that autophagy might responsible of BA mechanism of action.

Alternatively, some studies suggested that boric acid acts as histone deacetylase inhibitor (HDACi) [8] reduce intracellular Ca^++^ signals and storage [17], EIF2α/ATF4 and ATF6 Pathways [20, 21] or/and intervene in Wnt/β-catenin pathway [22].

In this study, we treated cancer cell lines from different organs that have a fully functional autophagy pathway. We found that low to medium concentrations of boric acid did not cause significant changes in cell growth in concordance with other studies, while effective doses were around 25mM. These doses are 700 to 3500 times higher than normal blood levels [13]. Although no reflection on health was observed when human population consumed water with Boron up to 29mg/L (equal to 2.5 mM boric acid) [23], nothing is known about concentrations as high as 25mM except that BA toxicity is not reported to be lethal [12].

After 72 hours of treatment with high doses of BA, a decrease in P62/SQSTM1 and accumulation of LC3B-II indicates a blockage in autophagy, but this happens after an initial increase in P62/SQSTM1 levels. The change of P62/SQSTM1 concentrations suggests a feed-back control that might be induced by other pathways, possibly due to BA effects on histone acetylation or mediating Ca^++^ signals. These concords with results from other studies that revealed changes in nucleic acid and transcription factors with BA treatments [21, 24].

We hypothesized that DU-145 is sensitive to boric acid due to the autophagy defect mentioned earlier, so we tried to reproduce autophagy arrest using 30µM of chloroquine. We established in our laboratory previously that the main autophagy pathway is inhibited in H1299 and COR-L23 cell lines at that concentration. Using chloroquine with and without small quantities of boric acid, we evaluated the response of our two NSCLC cell lines. In combination with 2mM of BA, the changes were minute. While at 5mM, the decrease in cell growth ranged between 7-10 %. The synergic effect is not strong enough (p>0.05). Increasing the boric acid concentration even further might increase the cellular response with chloroquine, but this contradicts the aim of decreasing BA concentrations to avoid possible toxicity. Thus, we do not see this combination of chloroquine and boric acid justified.

It was reported that other boron based compounds exhibit better anti-cancer properties such as Phenylboronic acid and Calcium Fructoborate [5, 7]. Those compounds presented a better effect which reflects in less toxicity, and future studies are recommended to focus on them.

Boric acid increases autophagy and causes dose-dependent cell death in cancers, and high concentrations can be therapeutical if real levels of toxicity were redefined above them. And although autophagy is modified with BA treatment, it is probably not the only player that induces cell death.

We recommend more investigation to understand boron role and its compounds and to get the most of this potential drug.

## 5. Acknowledgments

We thank Mrs Nieves Mateos for technical help and Hanna Jabbour for ideas and suggestions.

Funds for this project has been support via the European Commission (EU Partnerships and International Cooperation with Jordan, Lebanon, Syria and Palestine, EPIC projectgrant number is 2012-2624). This publication reflects the view only of the authors, and the Commission cannot be held responsible for any use that may be made of the information contained therein. Supported by ISC IIII-FEDER: PI13/01741.

## References

[1] T. A. Devirian and S. L. Volpe, “The Physiological Effects of Dietary Boron,” Critical Reviews in Food Science and Nutrition, vol. 43, no. 2, pp. 219–31, 2003.

[2] F. Nielsen, C. Hunt, L. Mullen and J. Hunt, “Effect of dietary boron on mineral, estrogen, and testosterone metabolism in postmenopausal women,” FASEB J, vol. 1, no. 5, pp. 394–7, 1987.

[3] W. T. Barranco and C. D. Eckhert, “Boric acid inhibits human prostate cancer cell proliferation,” Cancer Letters, vol. 216, p. 21–29, 2004.

[4] W. Barranco and C. Eckhert, “Cellular changes in boric acid-treated DU-145 prostate cancer cells,” British Journal of Cancer, vol. 94, p. 884–890, 2006.

[5] T. M. Bradke, C. Hall, S. W. Carper and G. E. Plopper, “Phenylboronic acid selectively inhibits human prostate and breast cancer cell migration and decreases viability,” Cell Adh Migr, vol. 2, no. 3, p. 153–160, 2008.

[6] E. M. McAuleya, T. A. Bradkea and G. E. Plopper, “Phenylboronic acid is a more potent inhibitor than boric acid of key signaling networks involved in cancer cell migration,” Cell Adhesion & Migration, vol. 5, no. 5, pp. 382–386, 2011.

[7] R. Scorei, R. Ciubar, C. M. Ciofrangeanu, V. Mitran, A. Cimpean and D. Iordachescu, “Comparative Effects of Boric Acid and Calcium Fructoborate on Breast Cancer Cells,” Biological Trace Element Research, vol. 122, no. 3, pp. 197–205, 2008.

[8] F. Di Renzo, G. Cappelletti, M. L. Broccia, E. Giavini and E. Menegola, “Boric acid inhibits embryonic histone deacetylases: A suggested mechanism to explain boric acid-related teratogenicity,” Toxicology and Applied Pharmacology, vol. 220, p. 178–185, 2007.

[9] D. Habes, S. Morakchi,. N. Aribi,. J.-P. Farine and N. Soltani, “Boric acid toxicity to the German cockroach, Blattella germanica: Alterations in midgut structure, and acetylcholinesterase and glutathione S-transferase activity,” Pesticide Biochemistry and Physiology, vol. 84, p. 17–24, 2006.

[10] J.-H. Huang, Z.-J. Cai, S.-X. Wen, p. Guo, X. Ye, G.-Z. Lin and L.-S. Chen, “Effects of boron toxicity on root and leaf anatomy in two Citrus species differing in boron tolerance,” Trees, vol. 28, no. 6, pp. 1653–1666, 2014.

[11] J. Zhanga, S. Nga, J. Wangc, J. Zhou, S.-H. Tan, N. Yang, Q. Lin, D. Xia and H.-M. Shen, “Histone deacetylase inhibitors induce autophagy through FOXO1-dependent pathways,” Autophagy, vol. 11, no. 4, pp. 629–642, 2015.

[12] A. o. t. s. a. d. r. (ATSDR), “Toxicological Profile for Boron,” Atlanta, GA: U.S. Department of Health and Human Services, Public Health Service., 2010. [Online]. Available: http://www.atsdr.cdc.gov/ToxProfiles/tp26.pdf. [Accessed 2016].

[13] K. Usuda, K. Kono and Y. Yoshida, “Serum boron concentration from inhabitants of an urban area in Japan. Reference value and interval for the health screening of boron exposure,” Biol Trace Elem Res, vol. 56, no. 2, pp. 167–78, 1997.

[14] S. Mahabir, M. Spitz, S. Barrera, Y. Dong, C. Eastham and M. Forman, “Dietary Boron and Hormone Replacement Therapy as Risk Factors for Lung Cancer in Women,” Am J Epidemiol, vol. 167, no. 9, p. 1070–1080, 2008.

[15] F. Murray, “A Human Health Risk Assessment of Boron (Boric Acid and Borax) in Drinking Water,” Regulatory Toxicology and Pharmacology, vol. 22, no. 3, pp. 221–230, 1995.

[16] I. o. M. (.P. o. Micronutrients, “Arsenic, Boron, Nickel, Silicon, and Vanadium,” in Dietary Reference Intakes for Vitamin A, Vitamin K, Arsenic, Boron, Chromium, Copper, Iodine, Iron, Manganese, Molybdenum, Nickel, Silicon, Vanadium, and Zinc, Washington (DC): National Academies Press (US), 2001.

[17] K. Henderson, S. L. J. Stella, S. Kobylewski and C. D. Eckhert, “Receptor activated Ca(2+) release is inhibited by boric acid in prostate cancer cells,” PLoS One, vol. 4, no. 6, p. e6009, 2009.

[18] W. Qiu and Y. Tian, “AB254. DU145: a naturally occurring cell with ATG5-independent alternative macroautophagy,” Transl Androl Urol, vol. 5, no. Suppl 1, p. AB254, 2016.

[19] D.-Y. Ouyang, L.-H. Xu, X.-H. He, Y.-T. Zhang, L.-H. Zeng, J.-Y. Cai and S. Ren, “Autophagy is differentially induced in prostate cancer LNCaP, DU145 and PC-3 cells via distinct splicing profiles of ATG5,” Autophagy, vol. 9, no. 1, p. 20–32, 2013.

[20] S. E. Kobylewski, K. A. Henderson, K. E. Yamada and C. D. Eckhert, “Activation of the EIF2α/ATF4 and ATF6 Pathways in DU-145 Cells by Boric Acid at the Concentration Reported in Men at the US Mean Boron Intake,” Biological Trace Element Research, vol. 176, no. 2, p. 278–293, 2017.

[21] K. A. Henderson, S. E. Kobylewski, K. E. Yamada and C. D. Eckhert, “Boric acid induces cytoplasmic stress granule formation, eIF2α phosphorylation, and ATF4 in prostate DU-145 cells,” Biometals, vol. 28, no. 1, pp. 133–41, 2015.

[22] A. Doğan, S. Demirci, H. Apdik, O. F. Bayrak, S. Gulluoglu, E. C. Tuysuz, O. Gusev, A. A. Rizvanov, E. Nikerel and F. Şahin, “A new hope for obesity management: Boron inhibits adipogenesis in progenitor cells through the Wnt/β-catenin pathway,” Metabolism, vol. 69, pp. 130–142, 2017.

[23] B. S. Sayli, E. Tüccar and A. H. Elhan, “An Assessment of Fertility in Boron-exposed Turkish Subpopulations,” Reproductive Toxicology, vol. 12, no. 3, p. 297–304, 1998.

[24] A. S. Acerbo and L. M. Miller, “Assessment of the chemical changes induced in human melanoma cells by boric acid treatment using infrared imaging,” Analyst, vol. 134, pp. 1669– 1674, 2009.

